# Switching metalloporphyrin binding specificity of a *b*-type cytochrome to fluorogenic zinc by design

**DOI:** 10.1101/832923

**Authors:** B. J. Bowen, A. R. McGarrity, J-Y. A. Szeto, C. R. Pudney, D. D. Jones

## Abstract

Metalloporphyrins play important roles in areas ranging from biology to nanoscience. Biology uses a narrow set of metal centres comprising mainly of iron and magnesium. Here, we convert metalloporphyrin specificity of cytochrome *b*_562_ from iron (haem) to fluorogenic zinc protoporphyrin IX (ZnPP). Through a computationally guided iterative design process, a variant with a near total preference for ZnPP was generated representing a switch in specificity. The new variant greatly enhanced (≥60 fold) the negligible aqueous fluorescence of free ZnPP *in vitro* and *in vivo*.

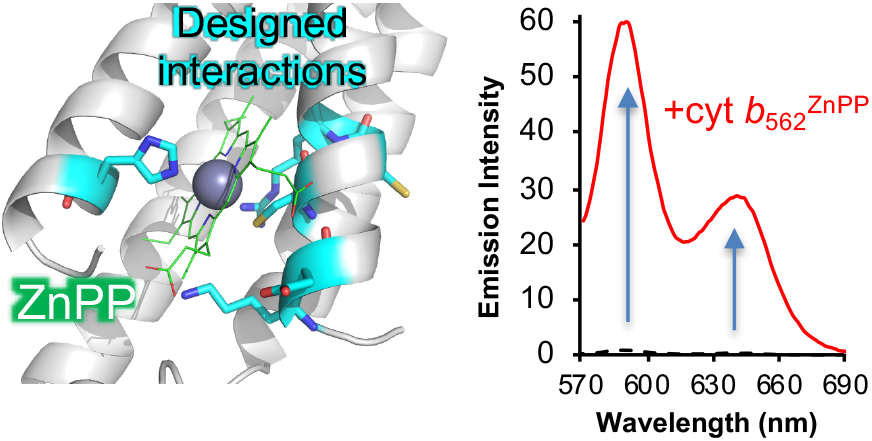

## Introduction

Here, we describe how an iterative design process can change the metalloporphyrin specificity of cytochrome *b*_562_ (cyt *b*_562_), a *b-*type haem binding protein unit, from iron to fluorogenic and photochemically active zinc. Metalloporphyrins ^1^ represent an important class of organometallic compounds and play a major role in biology as essential supplements to enable protein function in processes ranging from light harvesting to electron transport to enzyme catalysis to O_2_ transport ^2^. Binding to a protein tunes and modulates the metalloporphyrin physiochemical properties. The metal centres used in biology are Fe (e.g. haem), Mg (e.g. chlorophyll) and, to a much lesser extent Co (Vitamin B_12_) and Ni (F_430_). This does not however represent the full metal repertoire available to porphyrins (e.g. Cu, Pt, Ag, Cd, Ir, Zn) ^1^. There is currently great interest in incorporating new metal cofactors into proteins. Such abiotic cofactors have recently been shown to confer new enzymatic catalytic routes ^3–4^. Zinc porphyrins such as zinc protoporphyrin IX (ZnPP, Figure 1a) are one such class, playing important roles in clinical (including cancer therapy) and nanoscience settings. While ZnPP is not naturally utilised in biology as a protein cofactor, its aberrant presence in blood is commonly used to diagnose haem metabolism disorders ^5^. The photo-induced electron and energy transfer properties of zinc porphyrins make them particularly attractive for nanoscale applications, ranging from photosensitizers in solar cells to molecular optoelectronic components ^6–9^. ZnPP is inherently fluorescent but substantially quenched in aqueous solution ^10–11^. Designer protein components have the potential to further enhance ZnPP use through: (i) activation of fluorogenic properties ^12^ in aqueous conditions as an aid to clinical sensing and cell imaging; (ii) providing an organised, defined and tunable self-assembling scaffold for nanodevice construction ^6, 13–17^.

**Figure 1.**
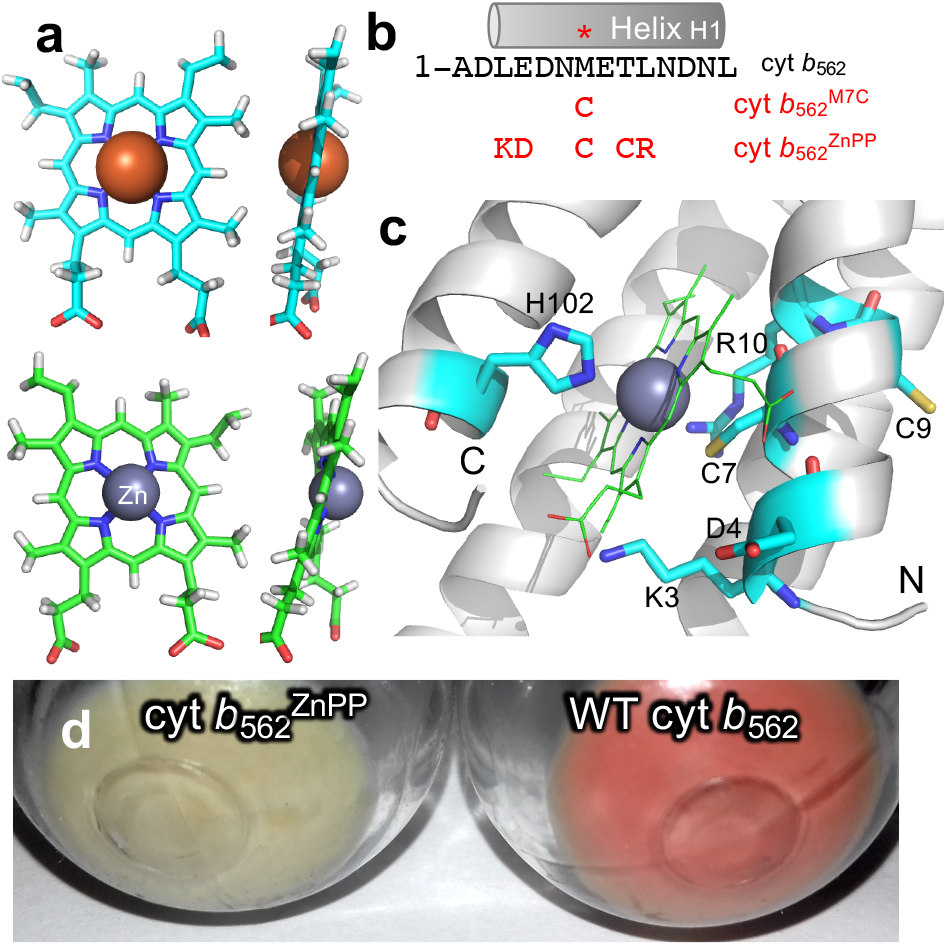
Porphyrin structure and cyt *b*_562_ variants. (a) Structure of haem (cyan) and ZnPP (green). The metalloporphyrin structure geometries were optimised using GAMESS-US ^21^. (b) Sequences of the cyt *b*_562_ variants and their corresponding nomenclature. The methionine that coordinate Fe in haem is highlighted with *. (c) Model of ZnPP-bound cyt *b*_562_^ZnPP^ with the residues mutated that shift metalloporphyrin specificity highlighted. (d) Cell pellets of recombinantly expressed cyt *b*_562_^ZnPP^ and the wild-type protein.

ZnPP incorporation within a protein scaffold has classically been achieved by either: (i) metal centre exchange (in the case of covalently attached porphyrins) that require harsh conditions unsuitable for most biological settings ^13, 18–19^; (ii) facile exchange in the case of non-covalent porphyrin-protein complexes ^11, 20^. With regards to the latter, affinity for ZnPP is inherently low (Table S1), with preference for the original haem cofactor; little attention is paid to optimising the protein interaction despite the structural and physicochemical variations on exchanging the porphyrin metal centre (Fig 1a). Here we show how computational design can be used to switch cyt *b*_562_ specificity from its normal iron porphyrin (haem) to zinc. The new protein can activate ZnPP fluorescence both *in vitro* and *in vivo*.

## Results and discussion

Cyt *b*_562_ (Figure S1) is a small, helical bundle protein that binds haem non-covalently ^22–24^. It is an important model for electron transfer ^25–28^, engineering novel components ^29–34^ and assembling supramolecular structures ^6, 14, 35^. ZnPP can replace haem through passive exchange ^20, 33, 36^ but the environment is non-ideal resulting in relatively low affinity binding (404 nM ±11 *K*_D_; Table S1) and a significant background haem binding *in vivo* (Figure 1). Factors include non-optimal metal coordination ligands and change in tetrapyrole structure and planarity due to the larger ionic radius of Zn^2+^(Fig 1a) ^37–38^.

The first stage in the design process was to optimise Zn coordination. Zn coordination is normally through soft ligands such as the cysteine thiol group and the imidazole group of histidine, or base ligands from the carboxylate groups of aspartate and glutamate; the native M7 *S*-methyl thioether group is not an ideal axial ligand. This was confirmed through computational analysis with cysteine considered the best, albeit still relatively poor, alternative (the M7C mutation). The second native axial ligand, the imidazole group of H102, was considered optimal so was not changed. The M7C variant was generated and the measured ZnPP and haem affinity (dissociation constant, *K*_D_) were comparable (181 ±11 nM for ZnPP and 206 ±2 nM for haem; Table S1) so generating a starting mutant with slightly improved ZnPP binding and lower haem affinity. Replacement of Met7 with either His (a second potential coordinating ligand) or Gly lowered ZnPP binding affinity to that of wild-type cyt *b*_562_ with the *bis*-His mutant retaining specificity for haem (Table S2). The absorbance spectrum of cyt *b*_562_^M7C^ (Fig S4) confirms ZnPP binding to the protein as indicated by the red shift in the Soret peak at 418nm for free ZnPP to a sharper peak at 431nm. The Soret and α/β band peaks of cyt *b*_562_^M7C^ are slightly blue shifted (8-10 nm) compared to the ZnPP bound to wild-type cyt *b*_562_. The Soret peak is also broader for wt cyt *b*_562_ suggesting a mixed binding population compared to cyt *b*_562_^M7C^. The ZnPP α/β peaks ratio also switches on cyt *b*_562_ binding.

To further optimise ZnPP binding, a more systematic *in silico* mutagenesis approach was adopted. Using the wt holo cyt *b*_562_ crystal structure 39 as a starting point, every residue from 1 to 20 was replaced to every possible residue and energy minimised to avoid clashes. The structures derived from the energy minimisation were used as the starting point for docking simulations using AutoDock ^40^ and Rosetta LigandDock ^41^. As a test for the accuracy, the *in silico* derived binding energy of the wt cyt was compared with the measured *K*_D_. The measured binding energy for wt cyt *b*_562_ (where ∆G=RT ln*K*_D_) is −8.7 kcalmol^−1^ for ZnPP, and −11 kcalmol^−1^ and for haem. These are close to the binding energies predicted from our modelling approach (Table S1). The *in silico* design process identified a variant with improved ZnPP binding energies termed cyt *b*_562_^ZnPP^. In addition to the M7C mutation, four other mutations were identified (L3K, E4D, T9C, L10R). The affinity of variant for ZnPP was much lower (<10 nM) compared to wt cyt *b*_562_ (Table S1 and Figure S3) with minimal haem binding observed both *in vivo* (Figure 1d) and *in vitro* (Figure S3a). This represents a near total switch in specificity from haem to ZnPP. The lowest scoring models were chosen for verification with Rosetta LigandDock, followed by molecular dynamics for 2 ns to ensure that the shape of the binding pocket was accurate. There was no significant change in the shape of the binding pocket when comparing before and after the molecular dynamics run. Residue 10 lies arose in the later stages of the design and buried in the holo form of the protein (Fig 1c) so it was surprising that not only was the L10R mutation tolerated but improved binding. Mutating residue 10 to either alanine or serine reduced affinity to >100 nM. Analysis of the model suggests that the guanidinium group of R10 lies within 2-3 Å of the C7 thiol group (Fig 1c and S4); proton abstraction by R10 may generate a thiolate group, which is a better ligand for Zn binding than the C7 thiol. Cyt *b*_562_^ZnPP^ was analysed in more detail.

Cyt *b*_562_^ZnPP^ had a characteristic absorbance spectrum indicative of formation of ZnPP holo protein (Fig 2a), with an extinction coefficient of 158 mM cm^−1^ at 424 nm, 5 nm blue shifted compared to wt cyt *b*_562_. On binding cyt *b*_562_^ZnPP^, ZnPP fluorescence intensity increased >60 fold (Fig 2b). Excitation at 431 nm gave the expected dual peak emission characteristic of ZnPP 19 (λ_EM_ 590 and 640 nm) with a Stoke’s shift of 160 nm and 210 nm (Fig 2b). The peak ratio of the two emission peaks was *circa* 2:1 (590:640 nm). Protein-bound ZnPP could be excited at 431 nm, 550nm and 590 nm (Fig S5). While the brightest emission originated from excitation at 431 nm the ability to excite at multiple and longer wavelengths has obvious benefits for biological imaging. These fluorescence characteristics were observed for wt cyt *b*_562_ and cyt *b*_562_^M7C^ (Fig S6) suggesting that the mutations affected only binding affinity. The quantum yield for the combined emission peaks was measured to be ~92% generating a protein with a brightness of 146 mM^−1^cm^−1^, higher than many autofluorescent proteins ^42–44^. ZnPP fluorescence is quenched in aqueous solutions to the extent that the quantum yield could not be accurately measured Coupled with the fluorogenic properties, cyt *b*_562_^ZnPP^ could make a useful genetically encoded imaging agent. A prerequisite for cell imaging is that ZnPP should pass through membranes and have no significant fluorescence in the cell until it binds specifically to cyt *b*_562_^ZnPP^. *E. coli* cells expressing cyt *b*_562_^ZnPP^ were colourless in comparison to those expressing wt cyt *b*_562_ (Figure 1d), which are a pink-red hue due to binding available haem with high affinity ^31^. This confirms cyt *b*_562_^ZnPP^ reduced affinity for haem and thus a decreased capacity to bind endogenous haem compared to the wt cyt *b*_562_. On addition of ZnPP to *E. coli* cells expressing cyt *b*_562_^ZnPP50^, fluorescence increased over time until plateauing after ~40-50 min (Fig 2c). Cells with no plasmid-encoded cyt *b*_562_ exhibited low baseline fluorescence on addition of ZnPP, which remained steady over the course of the incubation.

**Figure 2.**
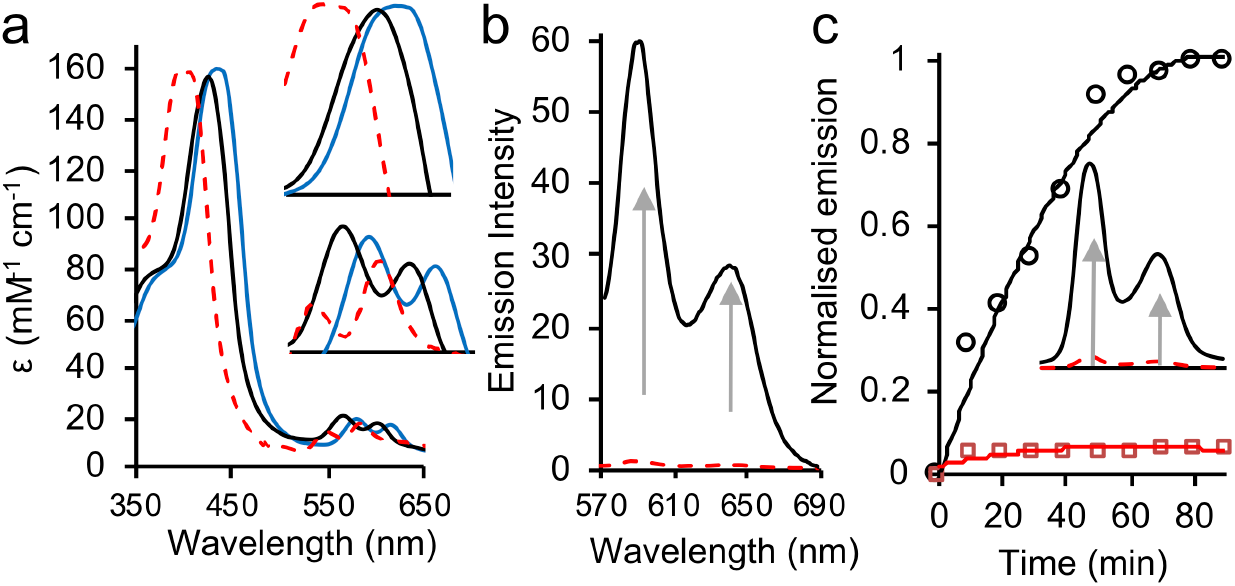
Absorbance and fluorescence properties of cyt *b*_562_ ZnPP binding variants. (a) absorbance spectra of free ZnPP (dashed red line), wt cyt *b*_562_ (blue line), and cyt *b*_562_^ZnPP^ (black line). Inset is the Soret peak (top) and α/β bands (bottom). (b) fluorescence emission spectra (on excitation at 431 nm) of ZnPP in the presence (black line) and absence (red dashed line) of cyt *b*_562_^ZnPP^. (c) Rate of fluorescence emission increase (excitation 431 nm) of *E. coli* cell culture incubated with ZnPP with cells containing (black line) or without (red line) plasmid-based cyt *b*_562_^ZnPP^. Inset is the fluorescence emission spectra after 1 hr.

Red edge excitation shift (REES) analysis was performed to ascertain the effect of ZnPP binding to cyt b_562_^ZnPP^ (Fig 3a). REES is an optical phenomenon where decreasing the excitation energy (increasing the excitation wavelength, *λ*_EX_) gives rise to a red shift in the maximum of the fluorescence emission (*λ*_EM_). Such shifts can arise where the fluorophore exists in a range of discrete solvation environments that are sampled as part of an equilibrium of solvent-solute interaction energies ^45^. In some cases, decreasing the energy of excitation allows for photo-selection of the discrete states that are red shifted (lower energy). We have recently demonstrated that for a single fluorophore (tryptophan) containing protein, the REES effect can inform on the equilibrium of conformational states accessible to the fluorophore ^46^. The REES data for free and protein-bound ZnPP are shown in Figure 3a and have been extracted for the α/β emission maximum shown in Figure 2b, *λ*_EM-1_ and *λ*_EM-2_, respectively. Free ZnPP exhibits a significant REES effect that is reduced on binding to cyt *b*_562_^ZnPP^; the change in *λ*_EM-1_ and *λ*_EM-2_ respectively being 0.56 ± 0.02 nm^−1^ and 0.21 ± 0.02 nm^−1^ for free ZnPP, which reduces to 0.12 ± 0.01 nm^−1^ and −0.04 ± 0.03 nm^−1^ on binding cyt *b*_562_^ZnPP50^. These data therefore suggest that the free ZnPP exists in an equilibrium of discrete solvation states and that this equilibrium collapses to essentially a single state on binding to the protein. These data therefore suggest tight ZnPP binding within the core of the protein, with limited solvent access, akin to haem binding. To further confirm ZnPP binding to cyt *b*_562_^ZnPP^, CD spectroscopy was used to monitor the expected transition from a partially folded helical structure to a folded 4-helix bundle on co-factor binding. On addition of ZnPP to cyt *b*_562_^ZnPP^, the deepening of troughs at ~208 nm and 222 nm was observed confirming the increase in helical character, with the spectra of holo-cyt *b*_562_^ZnPP50^ similar to that observed for wt holo-cyt *b*_562_ (Fig 3b).

**Figure 3.**
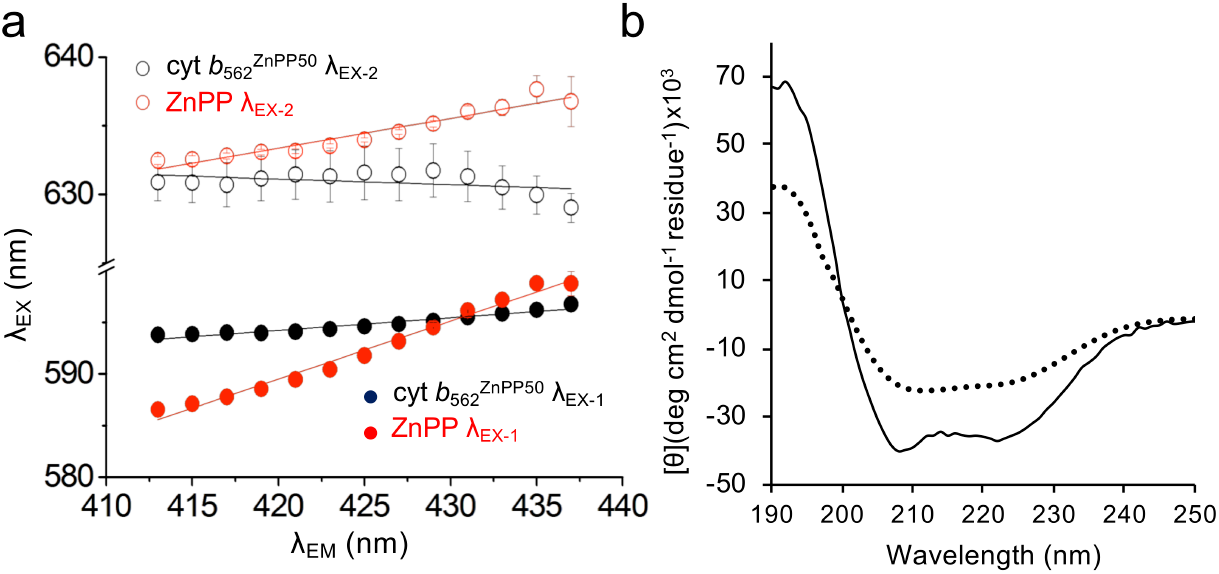
ZnPP binding to cyt *b*_562_^ZnPP^. (a) REES profile (emission wavelength, λ_EX_, *versus* excitation wavelength, λ_EM_) of free ZnPP (red line) and ZnPP-bound cyt *b*_562_^ZnPP50^ (black line). EX-1 and EX-2 refer to the two separate emission peaks. (b) CD spectra of apo-cyt *b*_562_^ZnPP50^ (black dotted line) and cyt *b*_562_^ZnPP50^ bound to ZnPP (black line).

Using systematic computationally guided engineering, the metalloporphyrin specificity of cyt *b*_562_ can be switched to ZnPP; the affinity is on a par with the natural haem cofactor and represented a near total switch in specificity. A fluorogenic effect was observed generating a protein complex with high brightness that exceeds that of many autofluorescent proteins. Higher affinity and thus more stable binding make holo-cyt *b*_562_^ZnPP50^ a more useful photo-electronic nanodevice ^15, 33^; a protein scaffold can also tune facets such as electronic excitation and emission. More generally, it provides a route for improving affinity of various different haemoproteins for ZnPP and other metalloporphyrins, in which the haem can be replaced by the photosensitizer effect of ZnPP ^13, 47^ or the catalytic properties of Ir and Cu porphyrins ^3–4^. The design process has shown changing metal coordinating ligands is not the defining step but that optimising residues adjacent to the metal coordination site is critical to success.

## Acknowledgements

DDJ and ARM would like to thank the Advanced Research Computing @ Cardiff facility, especially Thomas Green for help with access and usage of the Raven cluster. DDJ would like to thank the BBSRC (BB/H003746/1 and BB/ M000249/1), EPSRC (EP/J015318/1) for supporting this work. ARM was supported by a Cardiff School of Biosciences personal studentship. BJB was supported by a SWBio BBSRC DTP studentship.

## Supporting Information

### Supporting Methods

#### *In silico* design

##### ZincExplorer

*In silico* protein design was performed by initially using the native wild-type cyt *b*_*562*_ sequence with the ZincExplorer web server ^48^(http://protein.cau.edu.cn/ZincExplorer). Successive mutations were assessed by selecting existing residues, replacing them and resubmitting them to the server. The relative scores returned from the server were used to determine if a mutation was likely to improve Zinc protoporphyrin IX (ZnPP) binding (see Figure S1). The first design phase focused on replacing the M7 co-ordinating residue with alternatives to identify a better potential axial ligand residue. The second design round built on the best scoring variant from the first round introduced secondary mutations around residue 7.

##### Mutant Modelling

Using the published crystal structure of wild type cyt *b*_562_ (PDB:256B ^24^) as a starting point, *in silico* mutagenesis was performed using the mutagenesis application within MacPyMol ^49^ to change a single residue to another. This process was automated using shell scripts to allow for the creation of a library of structure files that replaced residue 1-20 with every other possible residue. Each initial model was used as the starting point and energy minimised to avoid any clashes caused by the mutation introduction. The starting structure was then placed within a triclinic box with dimension of 6.4×6.1×7.8 nm. This was populated using the Simple Point Charge (SPC) water model ^50^ to solvate the system to a total number of 16986 solvent molecules. The system was first energy minimised by performing 500 steps of steepest descent method followed by 500 steps of conjugant gradient method. The lowest energy state of the system was used as the starting conformation for the docking simulations.

##### Ligand Docking with AutoDock

Structure files for ZnPP were made using Avogadro ^51^. Geometry optimization calculations were performed using GAMESS-US ^21^ at the HF/6-31G* level in order to be consistent with the AMBER99sb force field ^52^. The optimised structure was used as the ligand file during docking simulations. The standard AutoDock4 ^40^ protocol was used as described in the software documentation. The scripts prepare_receptor4, prepare_ligand4, prepare_gpf4, prepare_dpf42 and prepare_flexreceptor4 (distributed with the software) were used to prepare the energy minimised mutated structure files for docking. For each variant 50 models were made and the lowest 5% of the predicted free binding energies were taken and averaged to give an approximate score for each mutation, these were then ranked and the lowest scoring mutant was chosen as the starting point for another round of mutations.

##### Ligand Docking with Rosetta LigandDock

The five lowest scoring variants from the AutoDock docking simulations for each round were chosen to be confirmed using the Rosetta LigandDock program ^41^. Docking was carried out using the Automatic RosettaLigand Setup script supplied with the program. The predicted binding energies and Rosetta score were compared with the results from the AutoDock docking analysis to inform the choice of which mutation would be used as the starting point of the next round of mutations.

#### Cyt *b*_562_ mutagenesis and protein production

Construction of cyt *b*_562_^ZnPP^ variants described in this paper was performed using site directed mutagenesis, essentially as outlined previously ^30^. Cyt *b*_562_ and its variants were expressed and purified as outlined previously ^30^.

When required, purified cyt *b*_562_ was converted to apo-form before ZnPP replacement by using organic solvent extraction approach ^31^. The proteins were buffer exchanged into water and the pH lowered to 1.5 before mixing with ice-cold 2-butanone. The mixture was then centrifuged for 2 minutes at 13,000x rpm in a bench top microfuge. The organic phase was removed and the preceding steps were repeated 5 times. The pH was returned to pH 6 before buffer exchanging back into 10mM Tris buffer.

ZnPP and hemin was purchased from Sigma Aldrich. The molecules were dissolved in 1 M NaOH prior to use to a final ZnPP concentration of 1 mM. ZnPP was stored in the dark at 4 C and centrifuged prior to use in experiments to prevent molecular aggregation and photodegradation. The ZnPP solution was then added to 10 mM Tris buffer to a final concentration of 200 μM. This was then added to a solution of 100 μM apo-cyt *b*_562_ in 10 mM Tris buffer and incubated for 12 hours at 25°C to form holo cyt *b*_562_ ZnPP.

#### Biophysical analysis of cyt *b*_562_ variants

##### UV-Vis Spectrophotometry and porphyrin titration

The absorbance spectra were monitored using Cary UV-Vis Spectrophotometer. The porphyrin titration experiments were performed using 20 μM apo-cyt *b*_562_ in 1ml of 10 mM phosphate buffer pH 6.2 with either 10 μM ferricyanide (for oxidised spectra) or 10 μM ascorbic acid (for reduced spectra). Either hemin (20 μM stock) or ZnPP (20 μM stock) were titrated against the protein solution. Apo-cyt *b*_562_ concentration was determined by measuring the absorbance at 280 nm (molar absorbance coefficient 2.98 mM^−1^ cm^−1^) and the holo-cyt b562 concentration was determined by monitoring 430 nm for ZnPP titration and 418 nm for oxidised hemin or 428 nm for reduced hemin. The resultant titration curves were then fitted to determine dissociation constant (see Figure S3 for examples), as described previously ^20^.

##### Fluorescence Spectroscopy

Excitation and emission spectra were recorded using a Varian Cary Eclipse spectrophotometer and the corresponding Cary Eclipse software. Samples of 750 μL comprising of 2.5-10 μΜ protein/porphyrin were analysed in a 5 × 5 mm QS quartz cuvette (Hellma, Müllheim, Germany). Spectra were recorded at a scan rate of either 120 or 600 nm/min with a slit width of 5 nm. Emission spectra were recorded up to 700 nm from a fixed excitation wavelength (corresponding to λ_MAX_ from the absorbance spectrum [~430 nm and ~590 nm] with an associated emission peak) for each variant. Quantum Yield studies were carried out using Alexa Fluor 431, essentially as described previously ^53–54^. For the *in vivo* analysis, liquid cell cultures were grown at 37°C to an A_600_ of 0.6. The cells were pelleted by centrifugation in a microfuge for 20 min at 4°C. ZnPP (20 μM) in 10 mM Tris-HCl (pH 8.0) was used to resuspended the cell pellet. A cell culture without the plasmid encoding the cyt *b*_562_ variants was used as a control. Fluorescence readings were taken periodically to determine rate of ZnPP uptake and incorporation.

##### Red edge excitation spectra (REES) analysis

All fluorescence measurements were performed using a Varian Cary Eclipse fluorescence spectrophotometer (Agilent Technologies) with a 5×5mm QS quartz cuvette (Hellma UK). The fluorimeter was baselined with 10mM sodium phosphate buffer (pH6.2) and all measurements were taken at 20°C. Excitation and emission slit widths were 5 nm and a scan rate of 600 or 120nm/min was used. Samples were incubated for 30 minutes at the given conditions prior to recording measurements. The red edge excitation scans were monitored from 413 to 437 nm with increments of 2 nm in the excitation wavelength. The wavelength of maximum emission, was extracted from the emission spectra by fitting to a sum (two) of skewed Gaussians of the form:

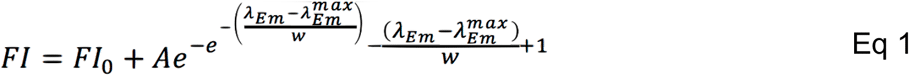

Where FI is the measured fluorescence intensity at an emission wavelength, λ_EM_. The spectra are then characterised by the amplitude, A of the spectrum at the maximum of fluorescence intensity, λ_EM_^max^, spectral width at half maximal, w and the minimum of the fluorescence emission, FI_0_, which was ~ zero for the buffer subtracted spectra.

##### CD spectroscopy

Circular dichroism spectroscopy was performed on a Jasco J-710 as described previously ^20^.

---

**Table S1.** Metalloporphyrin affinity of cyt *b*_562_ variants.

| Variant | Predicted Affinity (kcal/mol)<sup>a</sup> |  | Measured Affinity (nM) |  |
| --- | --- | --- | --- | --- |
|  | ZnPP | Haem | ZnPP | Haem |
| Wild Type | -8.7 | -12.3 | 404 | 10<sup>b</sup> |
| cyt *b*<sub>562</sub> M7C | -8.9 | -8.9 | 181 | 206 |
| cyt *b*<sub>562</sub><sup>ZnPP</sup> | -9.5 | -9 | <10<sup>c</sup> | ND<sup>d</sup> |

<sup>a</sup> calculated using Autodock. <sup>40</sup>; <sup>b</sup> determined previously <sup>20</sup>; <sup>c</sup> affinity too tight to measure accurately by current approach so a upper estimate is given; <sup>d</sup> affinity too weak to measure.

**Table S2.** Binding affinities for additional cyt *b*_562_ variants for ZnPP and haem.

| Variant | Measured binding affinity (nM) |  |
| --- | --- | --- |
|  | ZnPP | Haem |
| cyt *b*<sub>562</sub> M7H | 458 | 100 |
| cyt *b*<sub>562</sub> M7G | 409 | 472 |
| cyt *b*<sub>562</sub><sup>ZnPP</sup> L(R)10A | 152 | 200 |
| cyt *b*<sub>562</sub><sup>ZnPP</sup> L(R)10S | 108 | 248 |
| Cyt *b*<sub>562</sub><sup>ZnPP</sup> L(R)10C | 51 | 250 |

**Figure S1.**
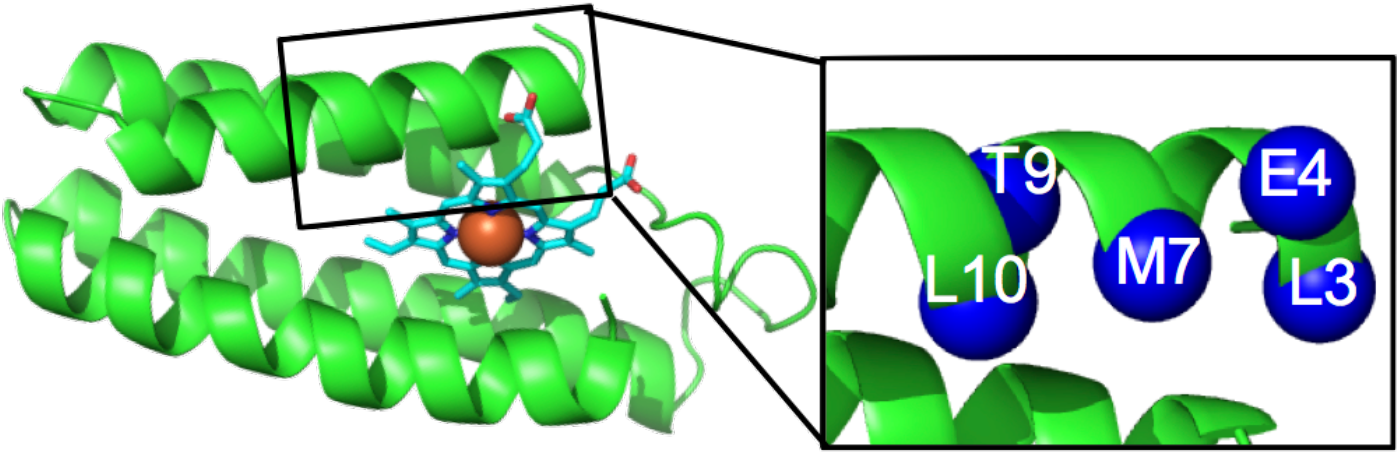
Cytochrome *b*_562_ structure. The protein is shown as green cartoon with haem shown in the stick representation and coloured cyan; the Fe centre is coloured brown. On the right hand side, the residues targeted for mutagenesis in helix H1 are shown.

**Fig S2.**
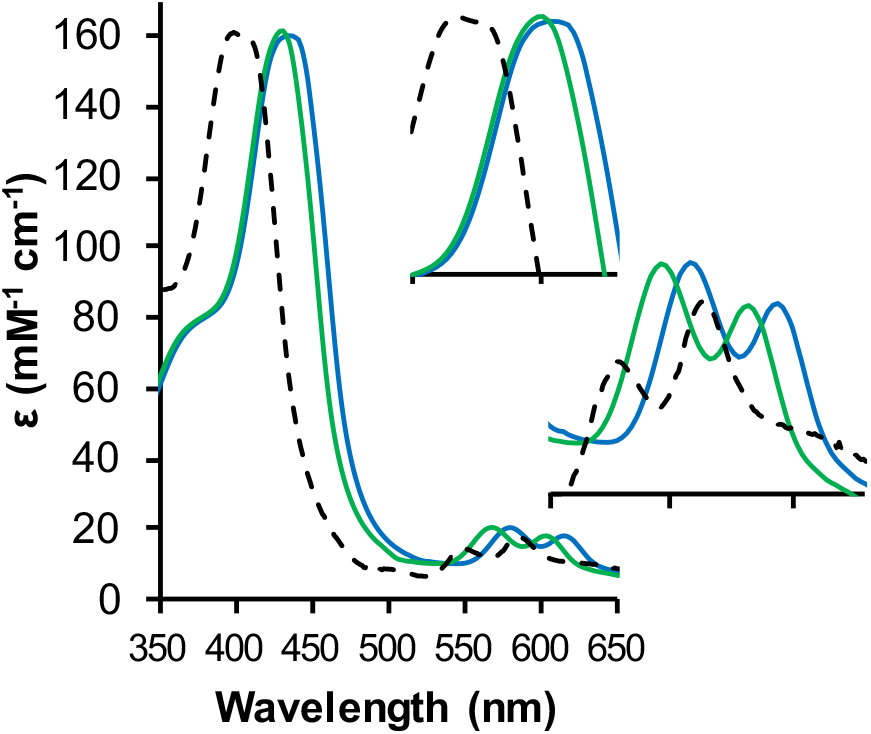
Absorbance spectra of free ZnPP (black dashed line) and bound to wt cyt *b*_562_ (blue line) and cyt *b*_562_^M7C^. Inset are enlarged images of the Soret peak (top) and α/β band (bottom).

**Fig S3.**
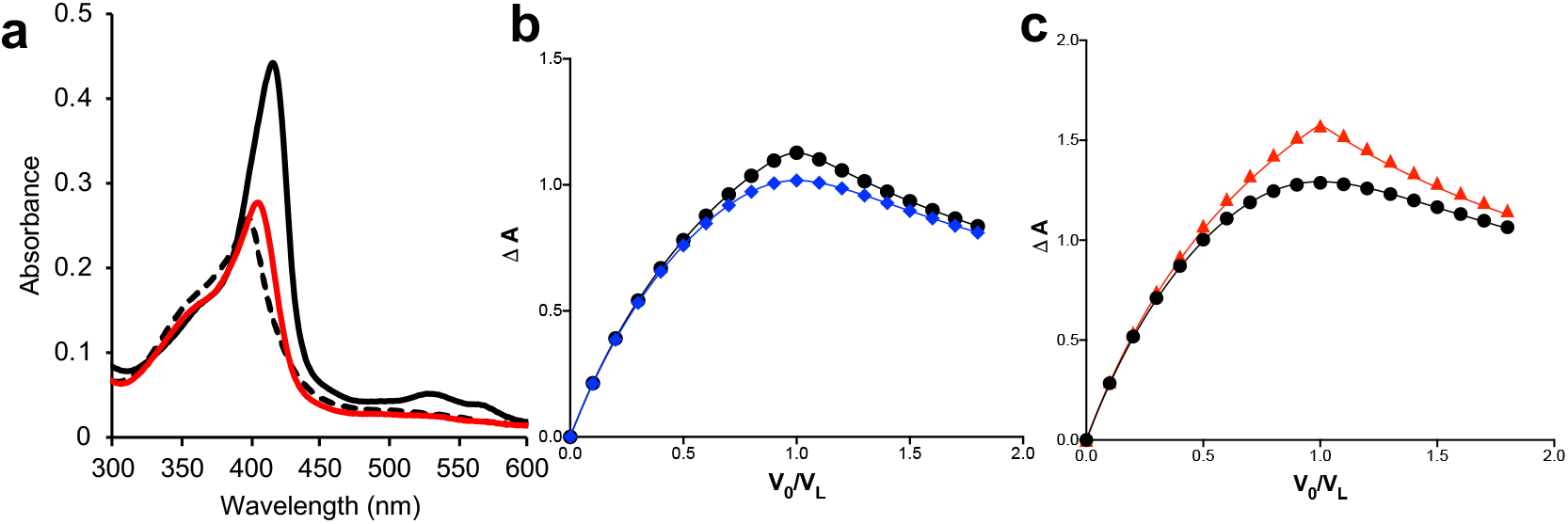
Binding affinity for haem and ZnPP. (a) cyt *b*_562_^ZnPP^ (red) and wt cyt *b*_562_ haem binding capacity. Protein (5 μM) was incubated with 6 μM haem and binding monitored. The free haem spectrum (dashed black lines) is shown for reference. Less than 10% haem binding to cyt *b*_562_^ZnPP^ was observed. (b) Haem and (c) ZnPP titration curves. The wild-type cyt *b*_562_ (black line, circles), cyt *b*_562_^M7C^ (blue line, diamonds) cyt *b*_562_^ZnPP^ (red) are shown. The data is based on triplicate reading (errors bars are removed for clarity). The data was fit and the *K*_D_ was calculated using a previously described approach ^20, 30^.

**Figure S4.**
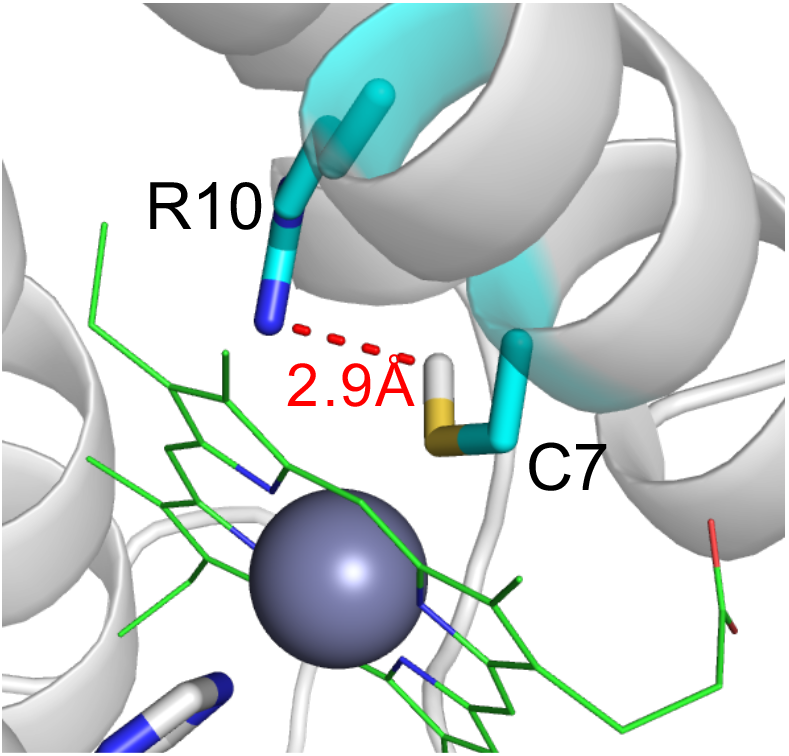
Model of the cyt *b*_562_^ZnPP50^ binding pocket. The potential interaction between the introduced guanidinium group of R10 and the new thiol group that can coordinate Zn^2+^ is shown as a red dashed line.

**Figure S5.**
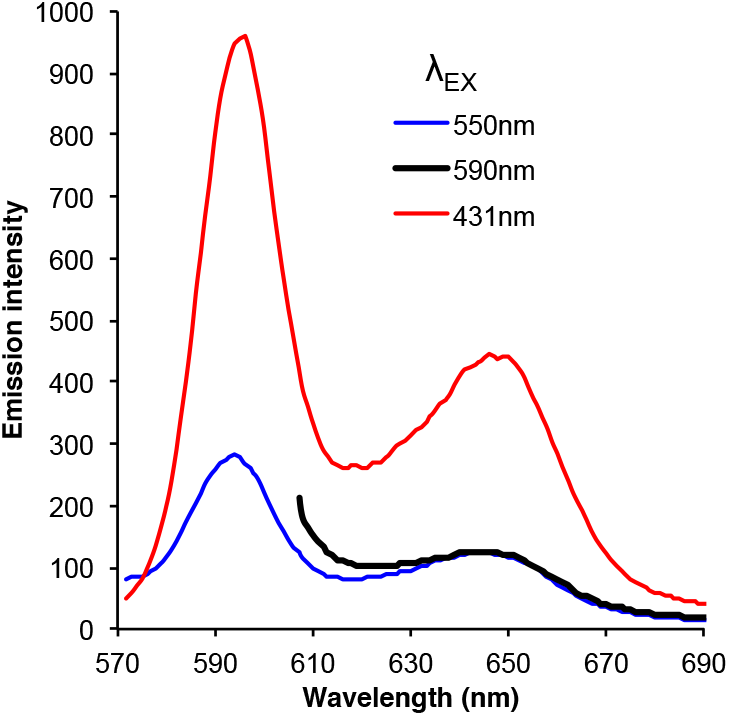
Relative fluorescence emission of profiles of ZnPP bound to cyt *b*_562_^ZnPP^ on excitation at the wavelengths stated in the figure.

**Figure S6.**
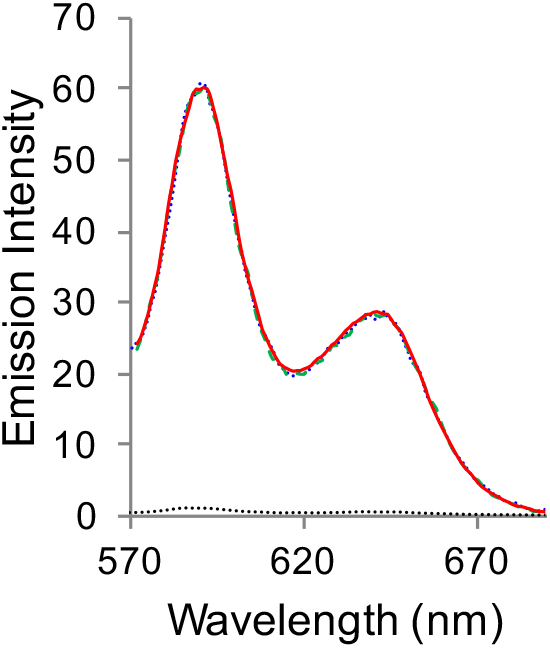
Fluorescence emission profile for cyt *b*_562_ variants bound to ZnPP. Red line, cyt *b*_562_^ZnPP^; dashed green line, cyt *b*_562_^M7C^; blue dotted line, wt cyt *b*_562_; dotted black line free ZnPP.

